# Tracking seasonal variation in modal properties of an open-grown green ash (*Fraxinus pennsylvanica*)

**DOI:** 10.1101/2025.10.06.680820

**Authors:** Daniel C. Burcham, Zuo Zhu, Siu-Kui Au

## Abstract

Despite longstanding interest and varied applications, the vibration behavior of trees remains poorly understood, largely due to methodological limitations. To more effectively identify and partition sources of natural variation in the modal properties of trees, there is a need for more reliable estimates derived from long-term observations. Using improved Bayesian methods, the modal properties of one green ash tree were identified over an entire growing season with simultaneous measurements of nearby weather conditions and phenological observations. The analysis confirmed many features of tree vibration observed in an earlier related study, including two prevalent close, nearly orthogonal modes and amplitude dependence for modal properties, but the more extensive and detailed measurements also revealed new characteristics, including changes in the arrangement of partial mode shapes and sizeable variation in modal properties over multiple time scales. For the leafless tree, the two modes were consistently oriented in directions aligned with the planting layout of surrounding trees, but their orientation varied erratically over time, while remaining roughly perpendicular to one another, during periods with leaves. Though some of the modal property variation closely paralleled other measurements, including weather conditions and vegetative phenophases, in physically reasonable ways, the reasons for other changes were not clear. Using a detailed assessment of the governing factors for estimate uncertainty, the cause of low-quality estimates was attributed to very close or nearly identical modes, and the causes and implications of close modes for tree vibration monitoring should be a main priority for future studies.

## 1. Introduction

Under normal conditions, trees dissipate kinetic energy absorbed from intermittent wind gusts by swaying at distinct frequencies (Gardiner, 1995; Schindler et al., 2013a; Schindler and Mohr, 2019), and the separation of wind excitation from energy dissipation processes reduces the likelihood of destructive harmonic resonance (Schindler and Mohr, 2019). For over 60 years (Sugden, 1962), forest scientists have investigated the vibration properties of trees using a variety of measurement and analysis techniques (Jackson et al., 2021). The studies routinely estimated modal frequencies, *f*_*i*_ (Hz), and damping ratios, *ζ*_*i*_ (dimensionless), for small numbers of trees at specific locations, and many reported dependable covariation between several intrinsic characteristics of trees, i.e., size (Jackson et al., 2021; Moore and Maguire, 2004), leaf condition (Baker, 1997; Bunce et al., 2019; Gougherty et al., 2018; Jaeger et al., 2022; Reiland et al., 2015; Schindler et al., 2013b), crown architecture (Miesbauer et al., 2014; Sellier and Fourcaud, 2005), and vibration properties across typical ranges (0.01 < *f*_*i*_ < 0.1; *ζ*_*i*_ << 1). While *f*_*i*_ was occasionally estimated using measurements of ambient tree vibration (Hassinen et al., 1998), *ζ*_*i*_ was only measured using free vibration tests in existing studies (Kane et al., 2014).

More recently, studies have shown similar covariation between other environmental conditions, especially air temperature (Bunce et al., 2019; Granucci et al., 2013) and water movement (Ciruzzi and Loheide, 2021, 2019; Kooreman, 2013; Raleigh et al., 2022), and tree vibration properties. Despite the strong covariance between tree vibration properties and several natural processes, it can be difficult to isolate changes associated with a specific process amid multiple sources of variability (Kooreman, 2013), especially during ambient vibration monitoring. Given the natural variability of tree modal properties, tracking and detecting changes in the structural properties of trees will be especially challenging, even compared to sophisticated methods developed for built structures (Yang et al., 2025). To effectively partition sources of variation, there is a need for more observations of the changes in tree modal properties over different time scales to improve analysis and modeling methods (Sellier and Suzuki, 2020).

The use of Bayesian inference for tracking the modal properties of trees was recently demonstrated using measurements of the ambient vibration of one large *Hopea odorata* (Dipterocarpaceae) over a two-week period (Burcham and Au, 2022). The analysis yielded hourly estimates of *f*_*i*_, *ζ*_*i*_, partial mode shapes, ***φ***_*i*_; and modal force PSD, *S*_*ii*_ (*g*^2^·Hz^−1^) with associated uncertainties represented using coefficients of variation, c.o.v. (%); and it revealed several novel features of tree vibration, including the prevalence of two close modes and amplitude dependence for *f*_*i*_ and *ζ*_*i*_ (Burcham and Au, 2022).

Though rare in most built structures, close modes with very similar frequencies usually occur in tall, slender structures with at least two transverse axes of symmetry, such as lighthouses (Brownjohn et al., 2019), tall buildings (Au et al., 2012), or even space launchers (De Vivo et al., 2013), and they are believed to arise from small deviations from axisymmetric conditions caused by design, construction, or minor defects. Compared to well-separated modes, close modes are much more difficult to identify because of the coherence between modal forces and much greater mathematical complexity. In the absence of similar existing reports, it is important to examine the occurrence of such novel features with more extensive observations, especially to identify potential challenges caused by close modes, and simultaneous measurements of wind flow, particularly to compare modal properties with wind conditions. Therefore, the objectives of this study were to:

1. characterize variation in the modal properties of one tree over an entire growing season,

2. examine amplitude dependence in modal properties using adjacent airflow measurements, and

3. examine mode alignment using adjacent airflow measurements.

## 2. Materials and Methods

### 2.1. Site and trees

One green ash (*Fraxinus pennsylvanica* ‘Sherwood Glen’) growing on the Colorado State University campus in Fort Collins, CO was selected for monitoring in the study (Figure 1). Approximately 25 years old at the time, the tree was 8.2 m tall with a trunk diameter at breast height (1.37 m above ground) of 17.8 cm. The tree was growing in a small stand of similar sized trees planted in a rectangular grid layout with 7.6 m between and 6.1 m within row spacing. The tree’s crown width along and across the planting row was 5.2 m and 4.5 m, respectively. During the growing season, the tree was irrigated regularly using two bubblers supplying 30 L of water on three separate days each week. The tree crown was not pruned or otherwise unnaturally altered before the study.

**Figure 1.**
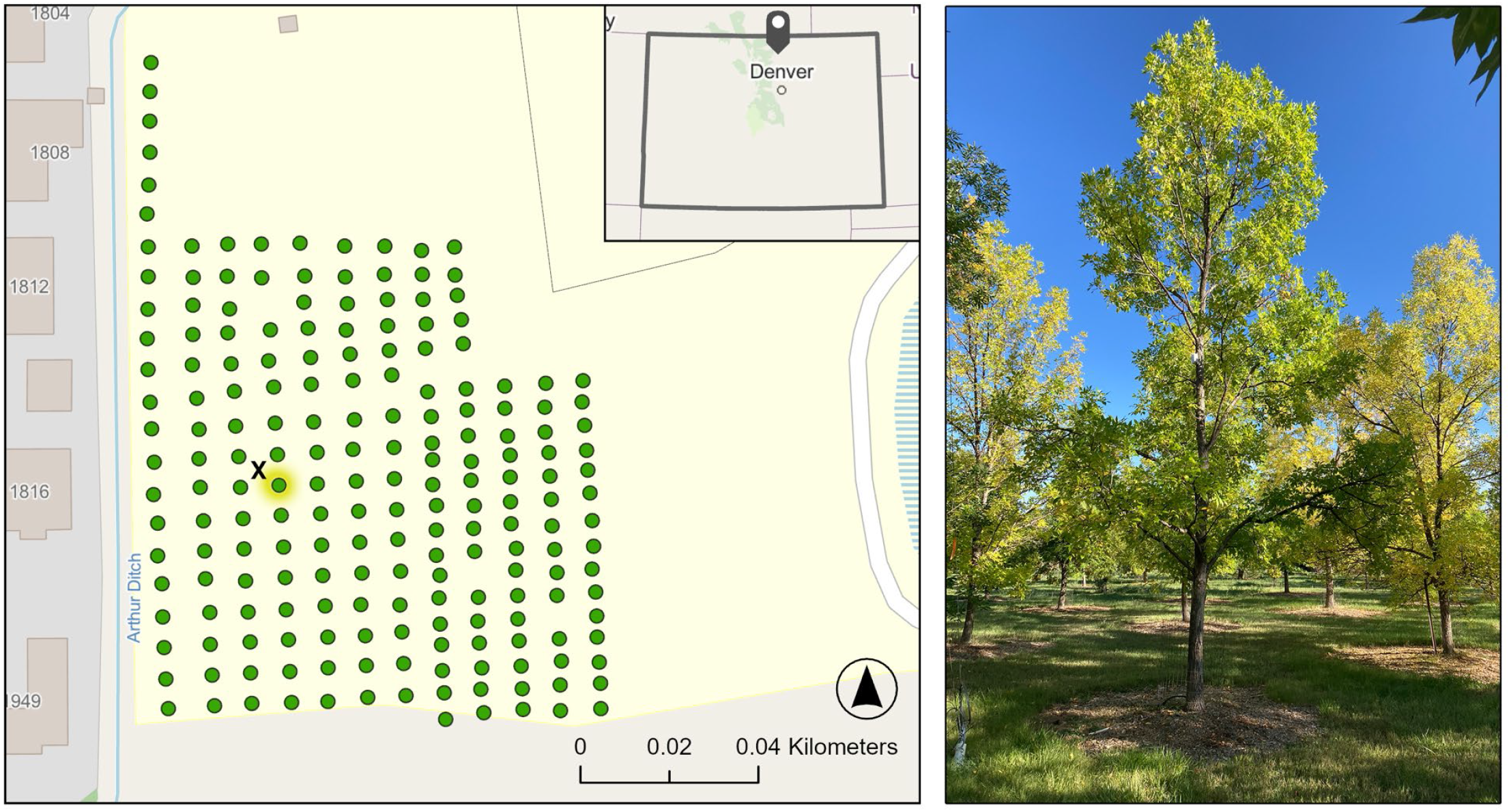
Using a triaxial accelerometer, the ambient vibration of an open-grown green ash (*Fraxinus pennsylvanica* ‘Sherwood Glen’, right) was monitored during the 2022 growing season. The tree (highlighted, left) was growing in the middle of a small stand of similar-sized trees (green markers, left) in Fort Collins, Colorado, USA (inset, left). During the study, wind conditions were also monitored using a triaxial ultrasonic anemometer mounted on a guyed mast (x marker, left) immediately next to the tree.

### 2.2 Measurements and data processing

To monitor tree movement, one triaxial accelerometer (AL100, Oregon Research Electronics) was installed 4.1 m above ground on the trunk. Using wood screws, the sensor was rigidly attached to the North side of the trunk to maintain the desired orientation and minimize solar heating of the electronics. The sensor’s *y*- and *z*-axes were oriented tangential (east-west) and perpendicular (north-south), respectively, to the underlying trunk surface, and the sensor’s *x*-axis was visually aligned parallel to the longitudinal axis of the trunk (approximately vertical). With a right-handed coordinate system, the sensor measured positive horizontal accelerations along the *y*- and *z*-axes towards the west and north, respectively. During operation, the accelerometer recorded tree movement continuously at 10 Hz.

To monitor wind conditions near the tree, a three-dimensional ultrasonic anemometer (81000, R.M. Young Co., Traverse City, MI, USA) was installed on a 12 m guyed sectional mast (PA2, South Midlands Communications, Chandler’s Ford, Eastleigh, UK). The mast was erected 8.2 m northwest of the ash, and the anemometer was mounted at 8.3 m, immediately above the canopy apex, on a 1 m boom extending to the south. It was expected that flow distortion from the guyed mast would be minimized by orienting the boom perpendicular to the prevailing wind direction. The anemometer’s *x*- and *y*-axes were oriented east-west and north-south, respectively, and its *z*-axis was oriented vertically. With a right-handed coordinate system, the anemometer measured positive wind velocities along the *x*-, *y*-, and *z*-axes from the east, north, and below, respectively. During operation, the anemometer measurements were continuously recorded at 20 Hz on a data logger (CR1000x, Campbell Scientific, Logan, UT) at 20 Hz.

To facilitate analysis, sixty-minute intervals starting at the beginning of every hour (i.e., hh:00:00.000) were used consistently for all signal processing and modal identification in this study. The duration was chosen to equally balance statistical accuracy (the longer the better) and conformance with modeling assumptions about stochastic stationary data (the shorter the better). For each interval used for analysis, local wind speed measurement outliers were identified as values more than three scaled median absolute deviations from the median and replaced with the nearest neighboring value. To account for possible vertical misalignment of the anemometer, the wind measurements were corrected using the planar fit method (Wilczak et al., 2001). Based on the assumption that streamlines parallel local terrain, the airflow measurements were rotated to align the anemometer’s *z*-axis perpendicular to the mean streamlines computed from 192 consecutive 30-minute intervals (i.e., four days). Subsequently, the wind measurements were rotated to align the anemometer’s positive *x*-axis with the prevailing (modal) wind direction. In the resulting streamwise coordinate system (Stull, 1988), the *u*- and *v*-axes were aligned with mean streamwise and cross-streamwise flow, respectively, and the mean speed along the *v*-axis was numerically equivalent to zero.

Using the streamwise wind measurements, two wind statistics were computed for each analysis interval, including the mean scalar horizontal wind speed, *Ū* (m×s^−1^):

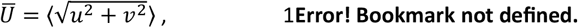

and kinematic momentum flux, *M* (m^2^×s^−2^):

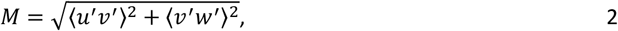

where *u*^′^, *v*^′^, and *w*^′^ are the fluctuations of the streamwise, cross-streamwise, and vertical wind speeds about their respective interval means following Reynold’s decomposition and ⟨·⟩ denotes averaging in the time domain.

Tree movement was monitored continuously between 15:00:00 on 4 April and 06:00:00 on 29 October 2022. Due to equipment malfunctions, wind conditions were monitored continuously for a shorter period between 00:00:00 on 21 April and 19:00:00 2 October 2022. To examine vibration properties associated with distinct stages of seasonal growth, the tree’s vegetative phenophases were monitored using standardized protocols (Denny et al., 2014; Denny and Crimmins, 2023) over the entire growing season. The start and end dates of five distinct phenophases were determined through regular visual observation, including breaking leaf buds, leaves, increasing leaf size, colored leaves, and falling leaves. For phenophase definitions, refer to the supplementary materials. During the transitional spring and fall seasons, the tree was observed twice per week to determine phenophases, but it was only observed once per week during the summer. Reproductive phenophases were not monitored because the tree’s expanding flower buds were damaged during a severe freeze (−10° C) on 13 April 2022.

### 2.3 Modal identification

Modal properties were identified using Bayesian operational modal analysis (BAYOMA). For a comprehensive introduction, see Au (2017) or Burcham and Au (2022) for a description of the method’s original application to trees. Using the Fast Fourier Transform (FFT) of ambient vibration data in specific bands modeled by classically damped structural dynamics, the method determines the posterior (i.e., given data) probability density function (PDF) of modal properties, including the natural frequencies; damping ratios; mode shapes; power spectral density (PSD) matrix of modal forces, ***S*** (g^2^·Hz^−1^); and PSD of noise, *S*_*e*_ (g^2^·Hz^−1^). With sufficient data, the posterior PDF can be approximated by a Gaussian PDF, which can be characterized in terms of the most probable value (MPV, the peak) and the remaining uncertainty (covariance matrix, the spread). The MPV is determined by minimizing a likelihood function with a specially designed algorithm for fast convergence (typically in a matter of seconds), and the covariance matrix is equal to the Hessian of the negative log-likelihood function evaluated at the MPV, also computed by special algorithms for speed and accuracy. In this work, a more recent algorithm that accelerates convergence based on Fisher Information Matrix scoring, called BAYOMA-FS, was used (Zhu et al., 2023). The Bayesian method requires two inputs, including a frequency band to define the FFT data used for identification and an initial guess of the natural frequencies, and they were determined by identifying resonance bands in the singular value spectrum (i.e., plot of eigenvalues of the PSD sample matrix against frequency) exhibiting dynamic amplification (Figure 2). To allow for variation in modal properties over time, the initial guess of frequencies was updated using a moving average of the MPVs over the preceding 24 intervals. The frequency band was similarly updated by setting the limits 0.1 Hz below and 0.25 Hz above the moving average of the 24 preceding MPVs of the modal frequencies. The uneven frequency band was used to exclude information associated with a lower secondary peak from the modal identification process while retaining a reasonably wide resonance band.

**Figure 2.**
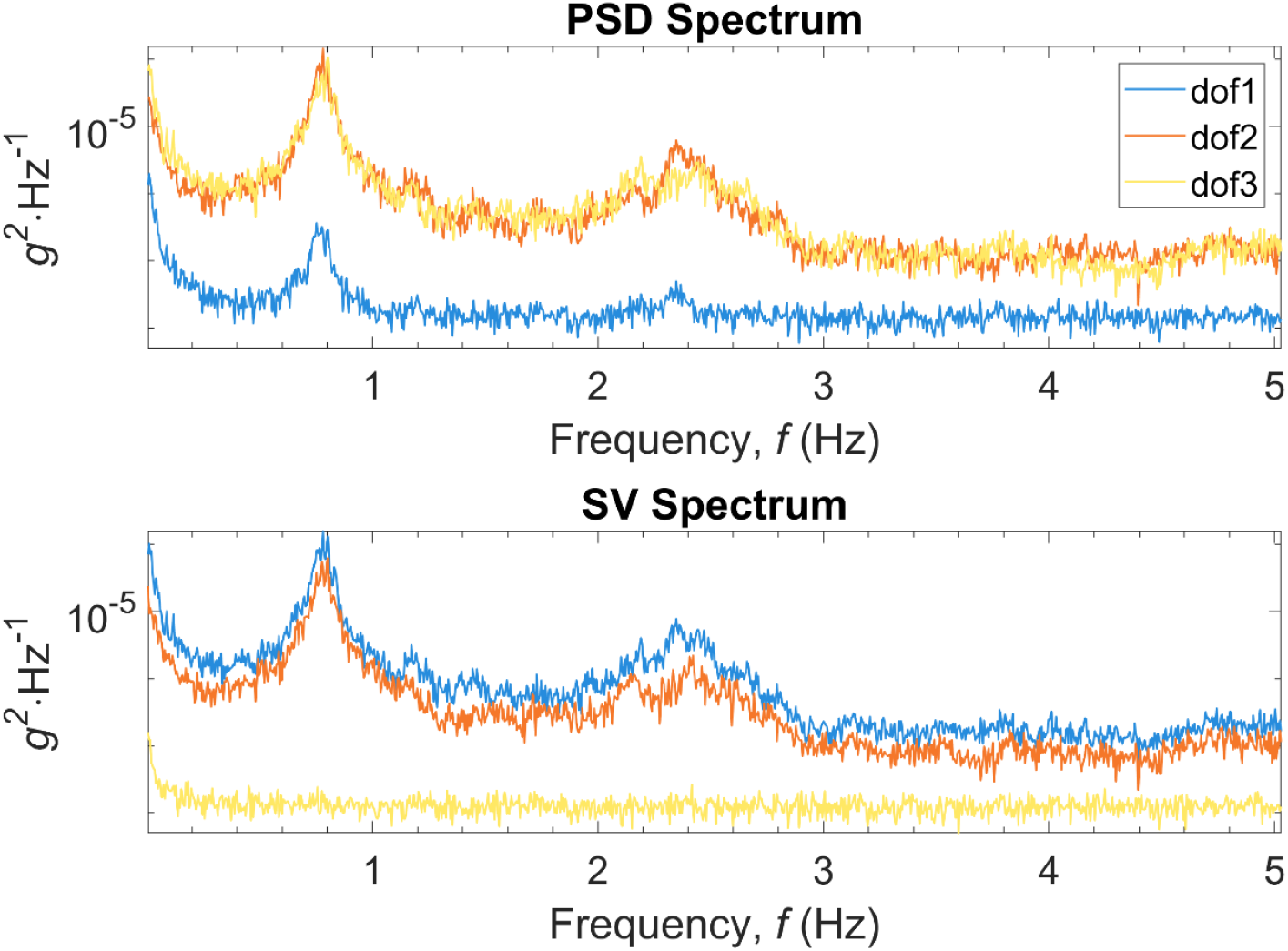
Power spectral density (PSD) and singular value (SV) spectra computed from one-hour time history of ambient vibration of a green ash (*Fraxinus pennsylvanica* ‘Sherwood Glen’) starting at 1500H on 4 April 2022. Note: dof1 = vertical; dof2 = radial (north-south); dof3 = tangential (east-west).

### 2.4 Data analysis

Tree modal properties were investigated after excluding poor quality estimates with signal to noise (s/n) ratios, defined as *γ* = *S*_*ii*_⁄4*S*_*e*_*ζ*^2^, below five and coefficients of variation (c.o.v.) for modal frequencies and damping ratios below 3% and 30%, respectively. The different c.o.v. limits were chosen because identification uncertainty for *f* is inherently much lower than that for *ζ*, with the two typically differing by at least one order of magnitude. To examine the relationship between weather conditions, phenophases, and modal properties, time histories of modal properties estimated from hourly intervals were examined alongside corresponding phenophase and weather records. The changes in modal properties were examined at both longer (e.g., seasonal) and shorter (e.g., daily) time scales. In addition to adjacent wind measurements, temperature and precipitation observations were acquired from a weather monitoring station located approximately 1.4 km north of the study site (Colorado Climate Center, 2022). During the study period, wet days were defined as the 24-hour periods following at least 3 mm of rain (Burns et al., 2015), and the classification was used to examine modal properties during periods with the expected additional mass from intercepted water (Ciruzzi and Loheide, 2021; Raleigh et al., 2022).

To examine possible amplitude dependence, *f*_*i*_ and *ζ*_*i*_ were plotted against modal force PSD, *S*_*ii*_, in bivariate scatter plots for periods with and without leaves separately. Due to the transitional nature of some phenophases, the intervening period after increasing leaf size and before falling leaves was used to examine the modal properties of the tree with leaves, and the period before breaking leaf buds and after falling leaves was used to examine the modal properties of the tree without leaves. Using the preceding definitions, distinct from the standardized phenophase definitions, amplitude dependence was not examined during periods of rapid change in leaf condition during the spring and fall. The relationship between wind conditions and *S*_*ii*_ was also examined by similarly plotting *Ū*and *M* against *S*_*ii*_ in bivariate scatter plots for periods with and without leaves separately.

The investigation of partial mode shapes, ***φ***_*i*_, a vector quantity, required some additional processing to reduce the number of dimensions for clarity and depict uncertainty correctly. Since the vertical component (*x*-axis) of ***φ***_*i*_ was negligible, the horizontal orientation of partial mode shapes was examined using angular directions. First, the actual orientation of ***φ***_*i*_ over time was depicted using a compass azimuth, representing the direction of each vector in terms of its clockwise rotation from the positive *z*-axis (aligned north). Second, the orientation of ***φ***_*i*_ with respect to the prevailing wind direction was examined in the streamwise coordinate system (defined previously) by determining the rotation of each vector from the positive streamwise (*u*+) axis. To determine the prevalence of orthogonality, the relative orientation of partial mode shapes was examined using the interior angle formed between the two vectors.

For dimensional consistency, the dominant mode shape uncertainties were converted to an angular uncertainty scaling with the representation of horizontal mode shape orientations. The dominant uncertainty of each mode shape can be obtained from the eigenvector ***u**_i_* (corresponding to the maximum eigenvalue) of its posterior covariance matrix scaled by the square root of the eigenvalue *δ*_*i*_ :

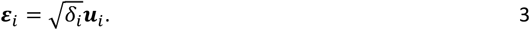

The dominant mode shape uncertainties, roughly tangential to the unit circle because the length of partial mode shapes (with a negligible vertical component) are constrained to one, were converted to an angular uncertainty defined as the sector between the most probable mode shape, ***φ***_*i*_, and ***ε***_*i*_ + ***φ***_*i*_.

Lastly, the posterior uncertainties for *f*_*i*_, *ζ*_*i*_, and ***φ***_*i*_ estimated using BAYOMA were compared with governing expressions for c.o.v.s, termed ‘uncertainty laws’ (Au et al., 2021; Au and Brownjohn, 2019). Analogous to the law of large numbers in statistics, the uncertainty laws are asymptotic formulas relating test configurations and structural properties to the expected identification uncertainty under specific conditions (i.e., long data, high signal-to-noise ratio, small damping, wide resonance band). Theoretically, the ratio of the posterior (BAYOMA) and expected uncertainties tends to one under situations consistent with modeling assumptions and accurate structural properties (Au et al., 2021). According to theory, the c.o.v.s of natural frequency and damping ratio are given by:

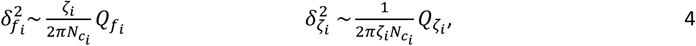

where 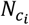 is a dimensionless data duration expressed as a multiple of natural period; 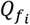and 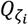 are coherence factors representing the amplification of uncertainty for close modes. The calculation of 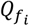 and 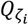 depend on multiple other factors, including modal entangling, modal disparity, and modal force coherence, in a complicated manner; see Au et al. (2021) for more information and all relevant expressions. The mode shape c.o.v. comprises contributions from two parts:

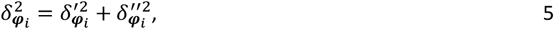

Where

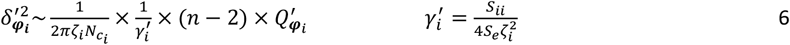

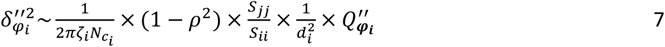

where 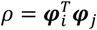 is the modal assurance criterion (MAC), *d*_*i*_ (dimensionless) is modal disparity (defined shortly), and 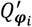 and 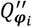 are the coherence factors for mode shapes (Au et al., 2021).

To explore potential causes of higher identification uncertainty, the influence of several factors governing uncertainties for close modes was examined in three separate plots. First, mode shape c.o.v. was plotted against modal disparity, *d*_*i*_ (dimensionless), a fundamental measure of the difference between two modes in terms of frequency and damping:

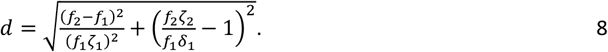

Second, the square root of the damping coherence factor, 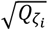, a measure of the amplification of damping c.o.v. caused by modal force coherence, *χ* (dimensionless):

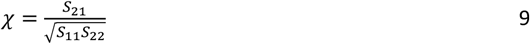

was plotted against the modulus of coherence, |*χ*|. Last, the absolute value of *ρ*, a measure of the proximity between ***φ***_1_ and ***φ***_2_, was also plotted against |*χ*|. All data processing and analysis was conducted using MATLAB R2024b (MathWorks, Natick, MA).

## 3. Results

During the spring season, the breaking leaf buds phenophase, with green leaf tips visible at the end of swollen vegetative buds, occurred between 29 April and 12 May, and the increasing leaf size phenophase started with the first completely unfolded leaf on 7 May and ended with most leaves attaining full size on 20 May. In the fall season, the colored leaves phenophase occurred between 27 September and 22 October, and the falling leaves phenophase started on 5 October and ended when no leaves remained on the tree on 22 October.

During the first hourly interval, the power spectral density and singular value spectra indicated the existence of two close modes near 0.74 and 0.79 Hz (Figure 2). Although two additional modes were evident near 2.4 Hz, modal properties were only identified for the two lowest modes. Using this information, the process consistently identified two similarly close modes over the entire growing season. Modal frequencies and damping ratios were separated by, on average, 2.7% and 2.2%, respectively. Comparatively, the algorithm converged more often during seasonal periods without (85%) than with (61%) leaves, despite higher *S*_*ii*_ for the latter. In total, the algorithm converged for 66% (3275/4975) of all hourly intervals, and, after excluding estimates with low s/n ratios and high uncertainties, 81% (2641/3275) of remaining hourly intervals were considered suitable for further analysis. For all modal properties, estimate uncertainty was lowest for *fi* with, on average, <1% COV over the entire season; the average uncertainty for *ζ*_i_ (11%) and *S*_*ii*_ (347%) was more than ten and 300 times greater, respectively. For all modal properties, estimate uncertainty was lower during periods without leaves, but the difference was greater for *S*_*ii*_ (457%) than *f*_*i*_ (0.4%) and *ζ*_*i*_ (0.6%) (Table 1).

**Table 1:**
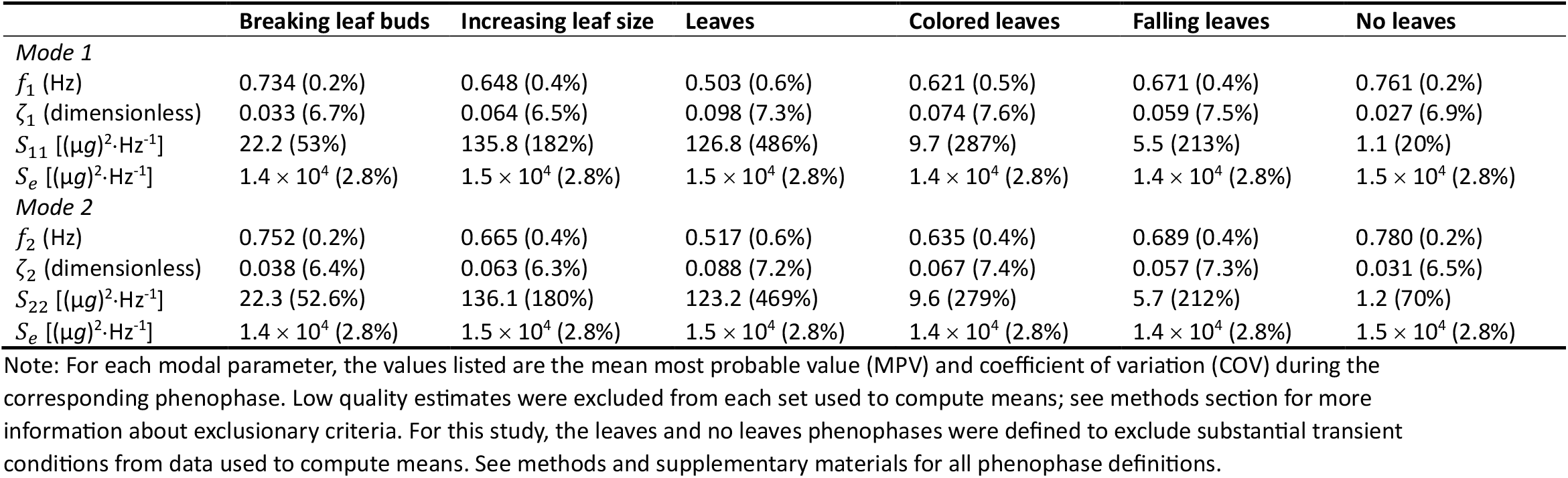
Summary of selected modal properties estimated during seasonal phenophases on an open-grown green ash (*Fraxinus pennsylvanica* ‘Sherwood Glen’)

|  | Breaking leaf buds | Increasing leaf size | Leaves | Colored leaves | Falling leaves | No leaves |
| --- | --- | --- | --- | --- | --- | --- |
| *Mode 1* |  |  |  |  |  |  |
| $f_1$ (Hz) | 0.734 (0.2%) | 0.648 (0.4%) | 0.503 (0.6%) | 0.621 (0.5%) | 0.671 (0.4%) | 0.761 (0.2%) |
| $\zeta_1$ (dimensionless) | 0.033 (6.7%) | 0.064 (6.5%) | 0.098 (7.3%) | 0.074 (7.6%) | 0.059 (7.5%) | 0.027 (6.9%) |
| $S_{11}$ [$(\mu g)^2 \cdot Hz^{-1}$] | 22.2 (53%) | 135.8 (182%) | 126.8 (486%) | 9.7 (287%) | 5.5 (213%) | 1.1 (20%) |
| $S_e$ [$(\mu g)^2 \cdot Hz^{-1}$] | $1.4 \times 10^4$ (2.8%) | $1.5 \times 10^4$ (2.8%) | $1.5 \times 10^4$ (2.8%) | $1.4 \times 10^4$ (2.8%) | $1.4 \times 10^4$ (2.8%) | $1.5 \times 10^4$ (2.8%) |
| *Mode 2* |  |  |  |  |  |  |
| $f_2$ (Hz) | 0.752 (0.2%) | 0.665 (0.4%) | 0.517 (0.6%) | 0.635 (0.4%) | 0.689 (0.4%) | 0.780 (0.2%) |
| $\zeta_2$ (dimensionless) | 0.038 (6.4%) | 0.063 (6.3%) | 0.088 (7.2%) | 0.067 (7.4%) | 0.057 (7.3%) | 0.031 (6.5%) |
| $S_{22}$ [$(\mu g)^2 \cdot Hz^{-1}$] | 22.3 (52.6%) | 136.1 (180%) | 123.2 (469%) | 9.6 (279%) | 5.7 (212%) | 1.2 (70%) |
| $S_e$ [$(\mu g)^2 \cdot Hz^{-1}$] | $1.4 \times 10^4$ (2.8%) | $1.5 \times 10^4$ (2.8%) | $1.5 \times 10^4$ (2.8%) | $1.4 \times 10^4$ (2.8%) | $1.4 \times 10^4$ (2.8%) | $1.5 \times 10^4$ (2.8%) |

Note: For each modal parameter, the values listed are the mean most probable value (MPV) and coefficient of variation (COV) during the corresponding phenophase. Low quality estimates were excluded from each set used to compute means; see methods section for more information about exclusionary criteria. For this study, the leaves and no leaves phenophases were defined to exclude substantial transient conditions from data used to compute means. See methods and supplementary materials for all phenophase definitions.

Over the entire monitoring period, large changes in the tree’s modal properties broadly corresponded to vegetative phenophases with distinctly lower frequencies, higher damping ratios, and higher modal force PSDs during periods with leaves (Figure 3). Compared to other quantities, the frequencies were more consistent across hourly intervals with smoother patterns of seasonal change. During the breaking buds and increasing leaf size phenophases, modal frequencies decreased 0.28 Hz over 21 days with most of the change occurring during the increasing leaf size phenophase. Over the summer, frequencies were generally stable around 0.5 Hz with a slight baseline trend towards higher values. During the colored and falling leaves phenophases, frequencies increased 0.21 Hz over 25 days with most of the change occurring during the falling leaves phenophase.

**Figure 3.**
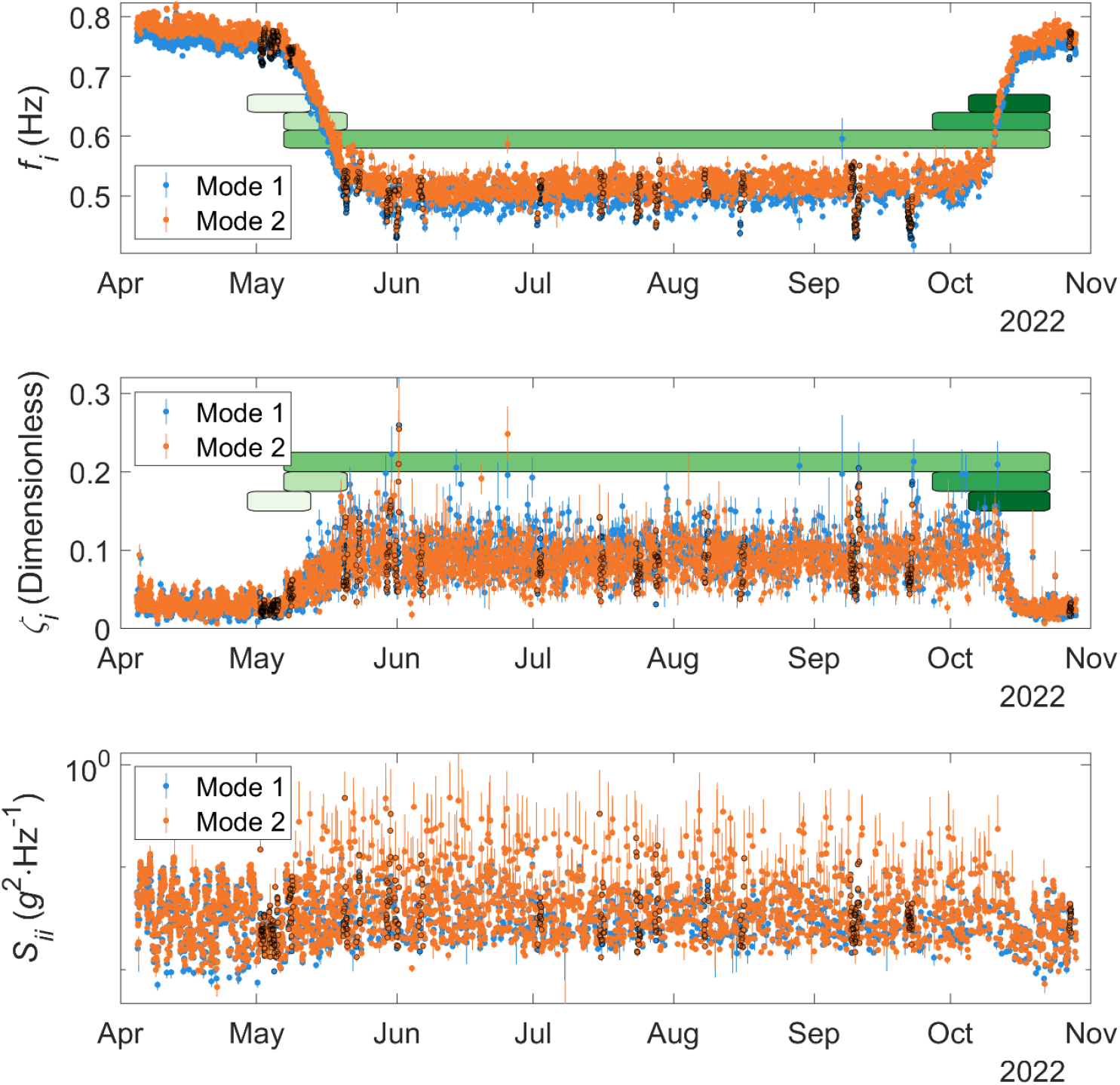
Modal properties (top-bottom: frequency, *f*_*i*_, damping ratio, *ζ*_*i*_, and modal force PSD, *S*_*ii*_) identified from hourly intervals of the ambient vibration of a green ash (*Fraxinus pennsylvanica* ‘Sherwood Glen’). Circle markers depict the most probable value (MPV) with a ±2*σ* error bar showing identification uncertainty for modes 1 (blue) and 2 (orange). Black outlines around selected markers denote wet days, defined as the 24 hours following at least 3 mm of precipitation. Colored bars show the occurrence of five distinct phenophases, including (light-dark green) breaking leaf buds, increasing leaf size, leaves, colored leaves, and falling leaves. Refer to the supplementary materials for phenophase definitions. Note: negative lower limits of some error bars were not displayed on the log-transformed axis for *S*_*ii*_.

Modal frequencies also uniquely varied over shorter time scales. There were several pronounced negative spikes in modal frequencies at specific times of the season (Figure 3). The deviations typically lasted several hours, and most of the negative spikes coincided with wet days. In contrast, a positive spike in modal frequencies coincided with a severe freeze in the early morning on 13 April 2022. During leafless periods, there were also sinusoidal patterns in modal frequencies recurring over daily intervals with maxima and minima during the early morning and afternoon hours, respectively, but the cyclical patterns were not as clear during periods with leaves (Figure 4). Similar features were not as clearly visible in time histories of *ζ*_*i*_ and *S*_*ii*_.

**Figure 4.**
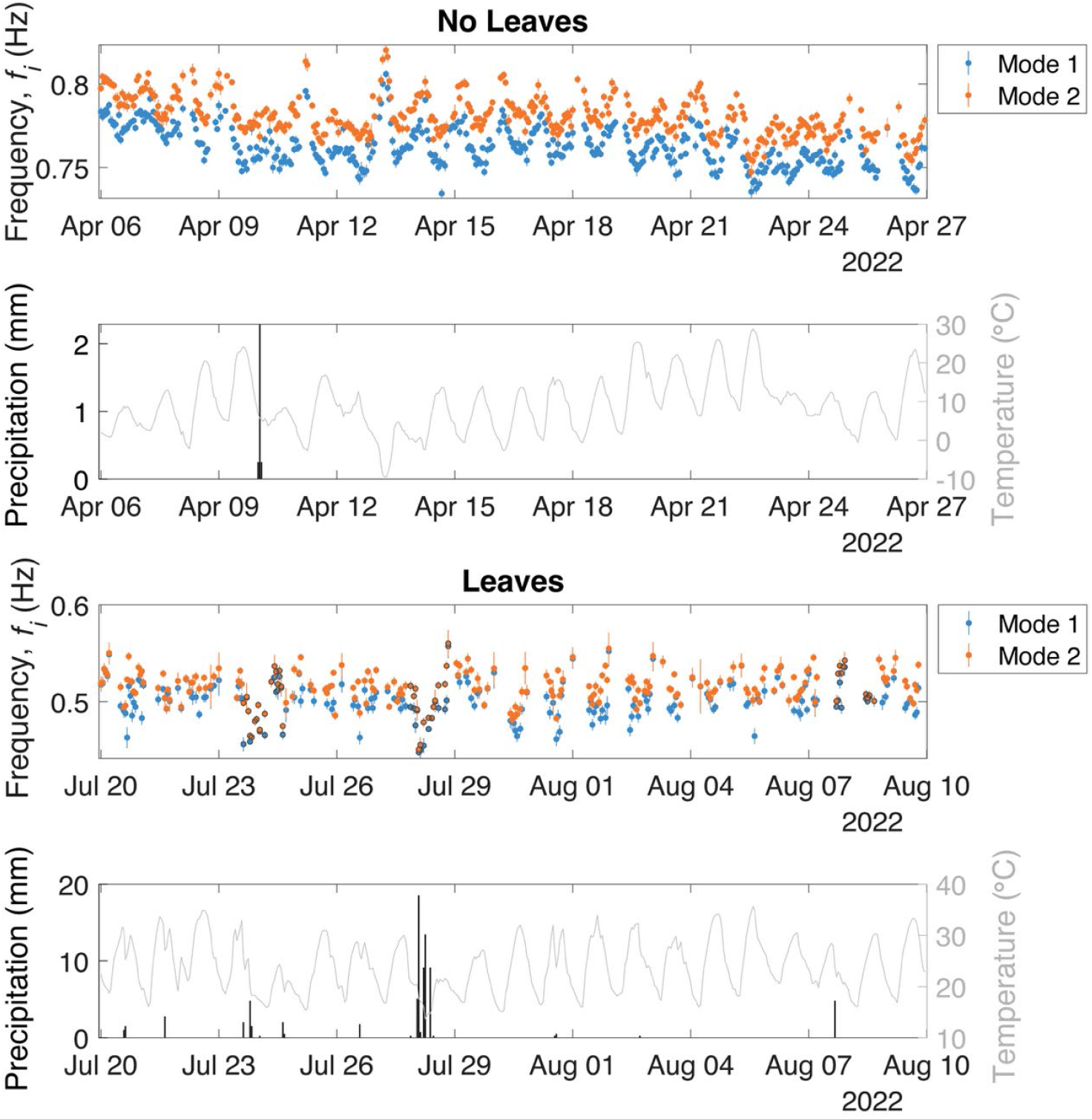
Modal frequencies identified from hourly intervals of the ambient vibration of a green ash (*Fraxinus pennsylvanica* ‘Sherwood Glen’) and corresponding hourly weather conditions, including total precipitation and mean temperature, for selected three-week periods without (top) and with (bottom) leaves. While diurnal variation in modal frequencies was evident during leafless periods (top), daily cycles were less visible during periods with leaves (bottom). In the early morning of 13 April, a positive deviation in modal frequencies (top) occurred alongside a mean hourly temperature close to −10° C. On multiple occasions, negative deviations in modal frequencies (bottom) corresponded with larger precipitation events. Black outlines around selected markers denote wet days, defined as the 24 hours following at least 3 mm of precipitation.

During the entire study, *f*_*i*_ and *ζ*_*i*_ covaried negatively and positively, respectively, with *S*_*ii*_, and the paired observations covaried more strongly, albeit with approximately twice the variance in modal properties, during periods with leaves (Figure 5). In contrast, the covariance between variables was weaker during periods without leaves. Over the observed range of *S*_*ii*_, for example, the range of *f*_*i*_ roughly doubled from about 0.5 to 1 Hz during periods with leaves compared to leafless periods. The covariance between modal properties and *S*_*ii*_ was consistently greater for mode 2, and the difference was exceptionally large between modal frequencies during periods with leaves. Similarly, wind conditions corresponded to a narrow and broad range of *S*_*ii*_ during periods without and with leaves, respectively (Figure 6). Comparatively, *S*_*ii*_ covaried more strongly with *M* than *Ū* during periods with and without leaves. During periods with leaves, *S*_*ii*_ was stratified across a wider range of values at a given *Ū* or *M*, and, for a subset of values, a given *Ū* or *M* corresponded to a much higher *S*_*ii*_.

**Figure 5.**
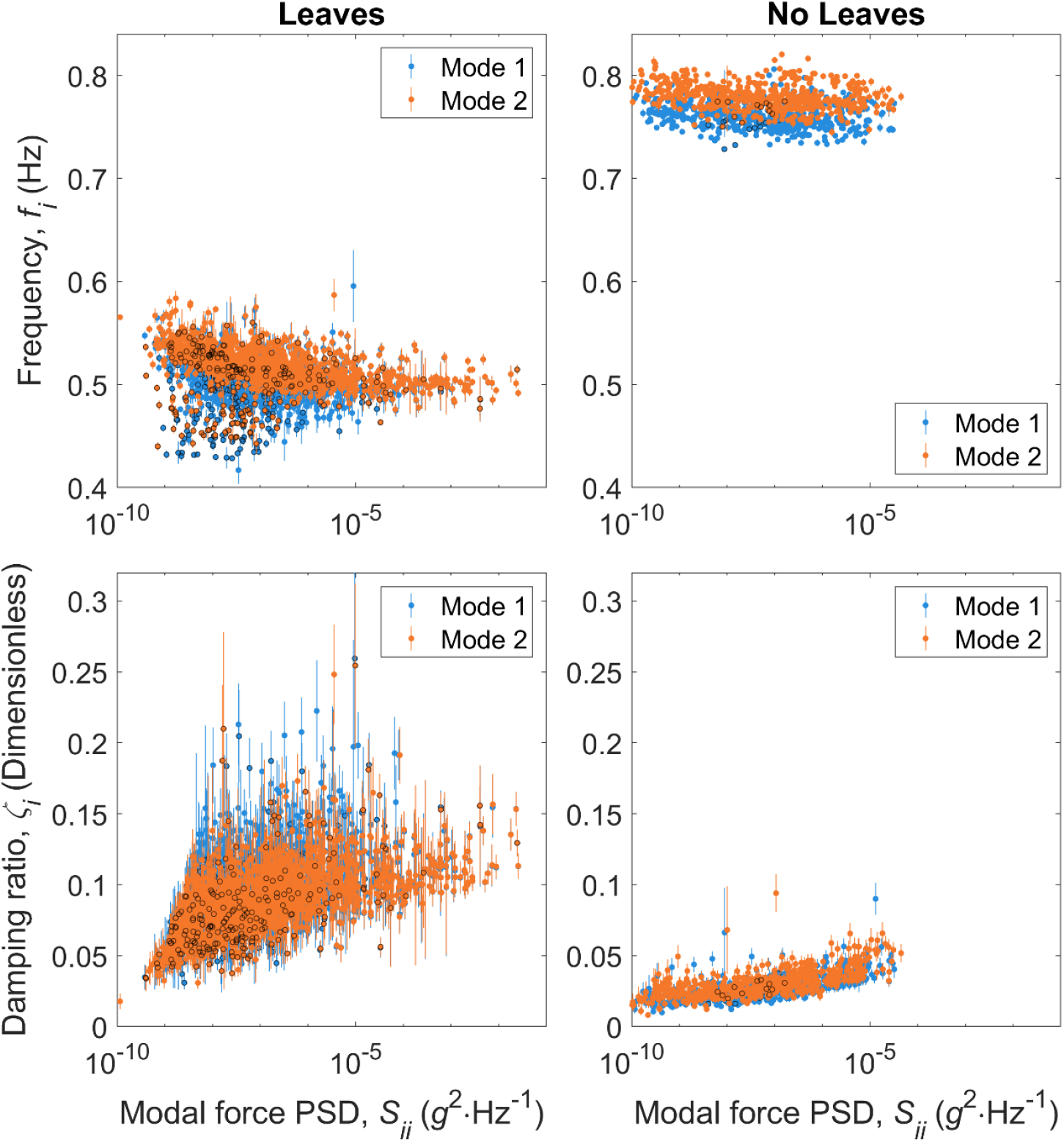
Scatter plots of modal frequencies, *f*_*i*_, (top) and damping ratios, *ζ*_*i*_, (bottom) against modal force PSD, *S*_*ii*_, during seasonal periods with (left) and without (right) leaves identified from the ambient vibration of a green ash (*Fraxinus pennsylvanica* ‘Sherwood Glen’). Circle markers depict the most probable value (MPV) with a ±2*σ* error bar showing identification uncertainty for modes 1 (blue) and 2 (orange). Note: black outlines around selected markers denote wet days, defined as the 24 hours following at least 3 mm of precipitation.

**Figure 6.**
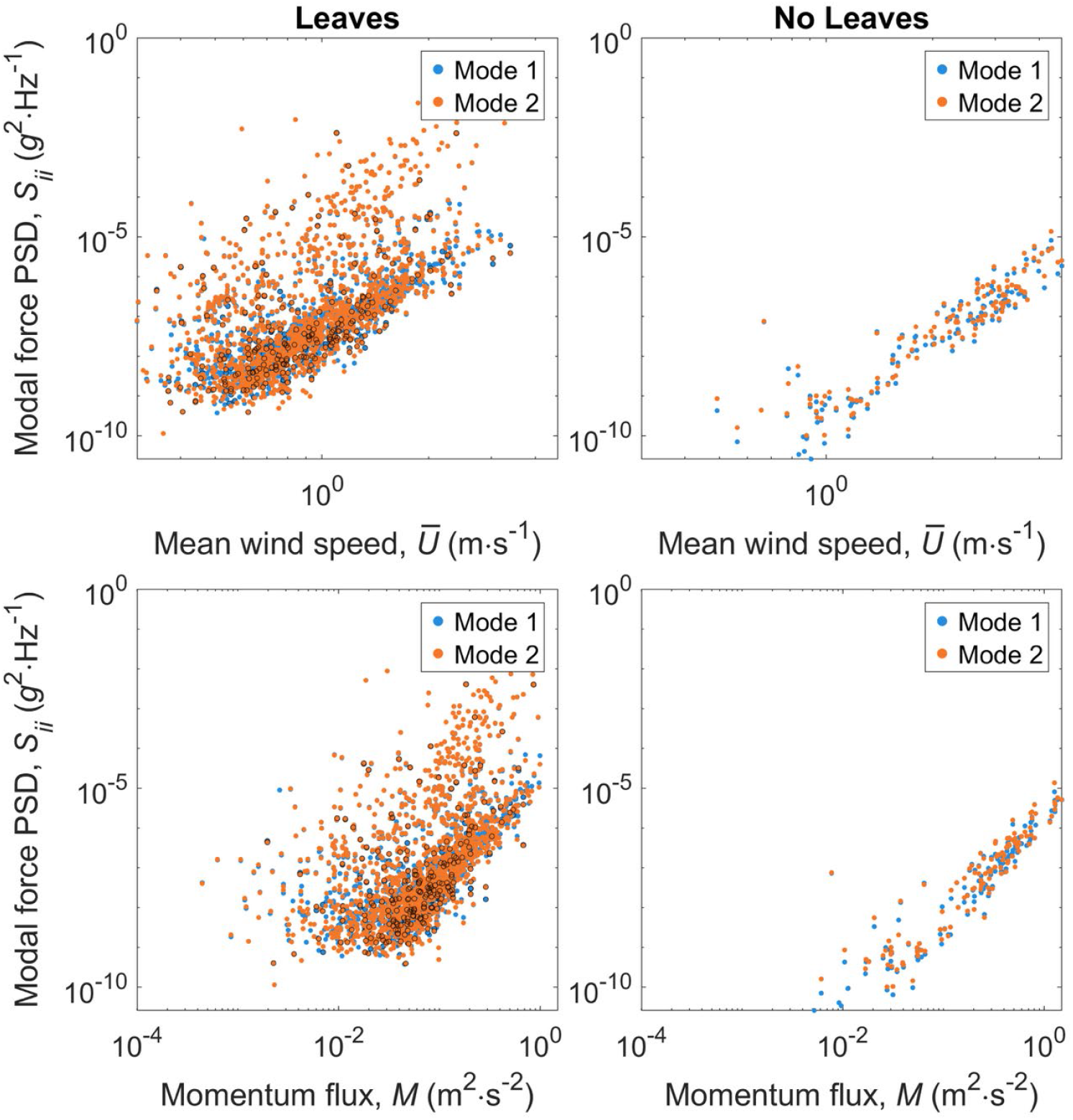
Scatter plots of modal force PSD against mean wind speed, *Ū* (m·s^−1^), (top) and momentum flux, *M* (m^−2^·s^−2^), (bottom) during seasonal periods with (left) and without (right) leaves identified from the ambient vibration of a green ash (*Fraxinus pennsylvanica* ‘Sherwood Glen’). Black outlines around selected markers denote wet days, defined as the 24 hours following at least 3 mm of precipitation.

During periods without leaves, ***φ***_1_ and ***φ***_2_ were more consistently oriented towards a limited number of directions (Figure 7) with ***φ***_1_ and ***φ***_2_ largely oriented across (east-west) and along (north-south) the planting row, respectively, in the stand of trees. During such periods, the modes were largely confined to two orientations along a single axis with ***φ***_1_ and ***φ***_2_ more frequently oriented towards the east and north, respectively. In contrast, the partial mode shapes were not oriented consistently over time during periods with leaves. Notably, the interior angle between ***φ***_1_ and ***φ***_2_ was regularly close to 90° during the entire study, but the partial mode shapes were occasionally oriented in the same (0° interior angle) or opposite (180° interior angle) directions during seasonal periods with leaves. In the streamwise coordinate system, the orientation of partial mode shapes showed a similar difference between seasonal periods without and with leaves. During leafless periods, ***φ***_1_ and ***φ***_2_ were aligned near the *v* (cross-streamwise) and *u* (streamwise) axes, respectively, with a slightly greater tendency to align towards the positive (downstream) axis directions. Regardless of the coordinate system, ***φ***_*i*_ c.o.v.s were much greater during periods with leaves. During the entire study period, the alignment of partial mode shapes appeared slightly more concentrated along the positive streamwise (*u*+) direction, especially for ***φ***_2_.

**Figure 7.**
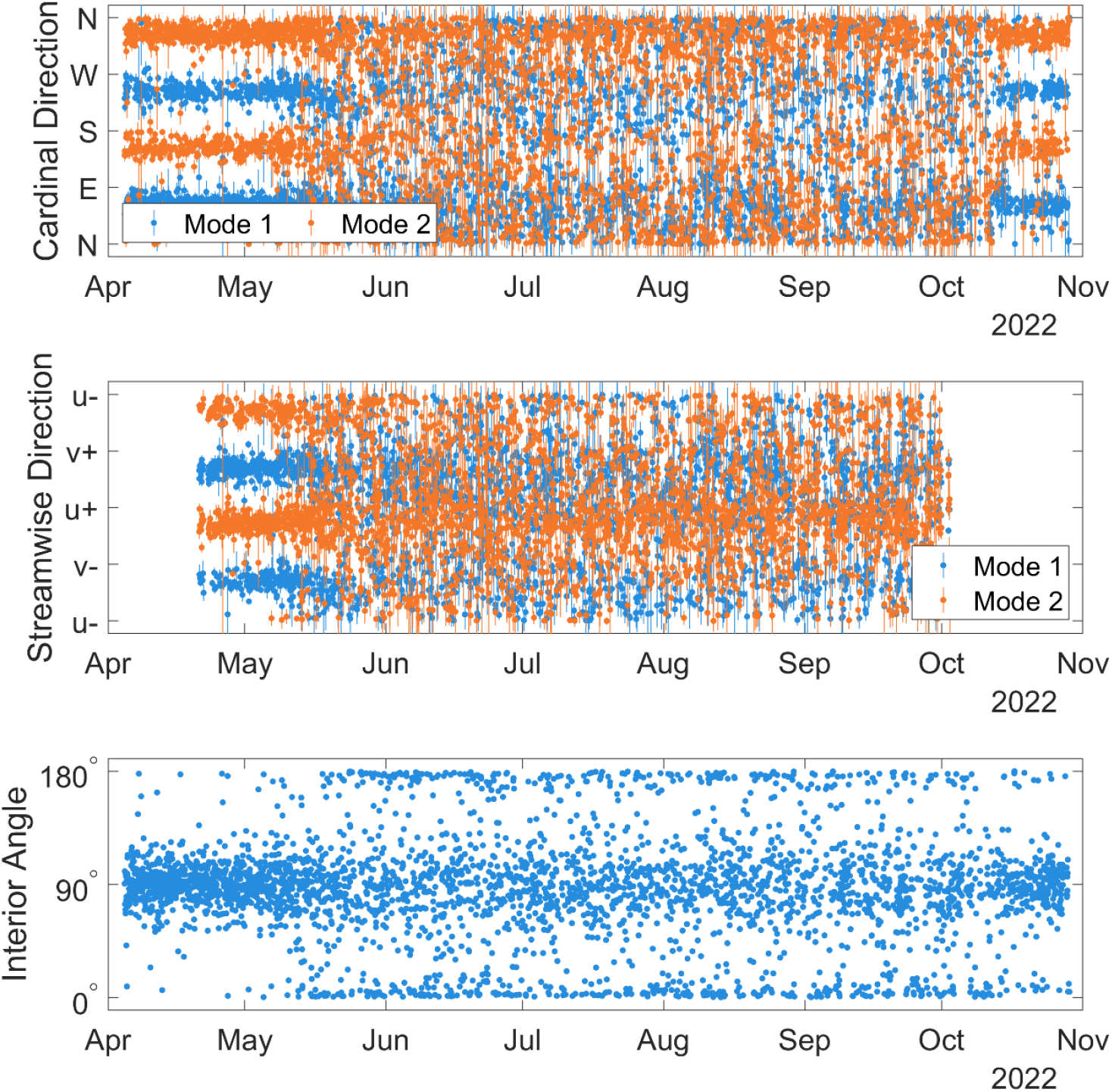
Horizontal orientation of partial mode shapes, ***φ***_*i*_, displayed as the clockwise rotation from north, i.e., compass azimuth, (top); clockwise, i.e., negative, or anti-clockwise, i.e., positive, rotation from the prevailing (downstream) wind direction (middle); and interior angle between the two vectors (bottom). For the two upper panels, circle markers depict the horizontal orientation of the most probable value (MPV) with a ±2*σ* error bar showing identification uncertainty for modes 1 (blue) and 2 (orange).

For most estimates concentrated near the lower limit of uncertainties, the posterior c.o.v. obtained from BAYOMA was similar to the values obtained from uncertainty laws with paired observations falling near a 1:1 line (Figure 8). However, there was a small portion of cases, approximately 20% of the total, with c.o.v.s predicted from uncertainty laws exceeding those from BAYOMA, and the cases generally corresponded to instances with either 0° or 180° between ***φ***_*i*_ (Figure 7). Closer inspection of the cases showing disagreement revealed either a low s/n ratio or very close modes. The examination of governing factors attributed the cause of such cases to very close modes with mode shape c.o.v. increasing for progressively lower *d*_*i*_ and both 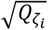 and *ρ* increasing with |*χ*|.

**Figure 8.**
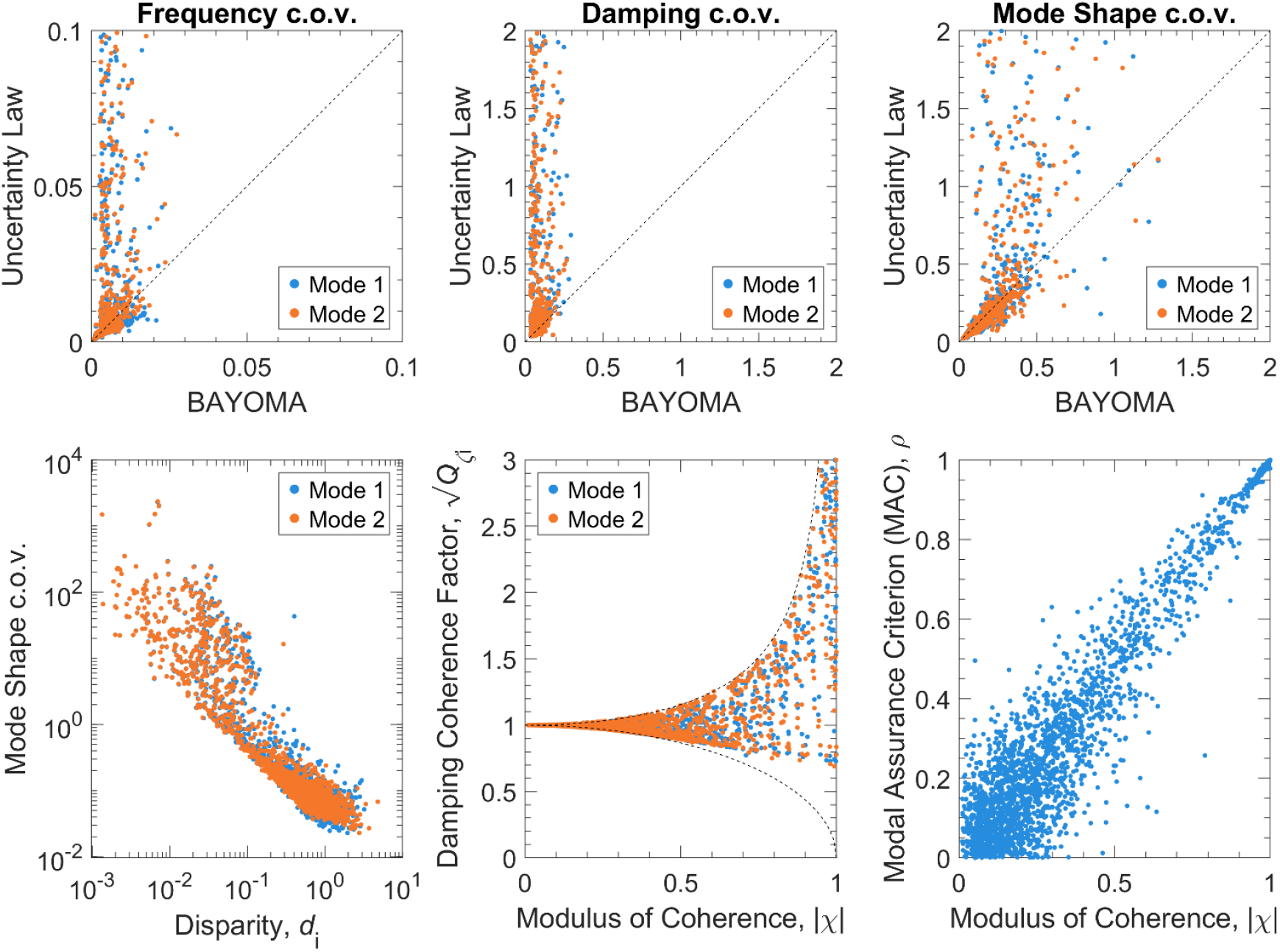
Summary of observed and expected uncertainties for all modal properties and selected governing factors. For modal frequencies, damping ratios, and partial mode shapes, the comparison of c.o.v. from uncertainty laws with the same from BAYOMA showed much lower uncertainties for the former in approximately 20% of all cases (top row). Showing one cause of higher uncertainties, the mode shape c.o.v. plotted against modal disparity (bottom left) showed an increase in uncertainty for close modes, especially for *d*_*i*_ < 0.1. Likewise for the damping coherence factor (bottom center) and MAC (bottom right) plotted against the modulus of coherence, higher modal force coherence with very similar modes, i.e., *ρ* ≈ 1, accompanied increased uncertainty for damping ratios.

## 4. Discussion

The modal properties identified using ambient vibration of the green ash were comparable to existing measurements of small broadleaf trees (Kane and James, 2011), and the broad pattern of seasonal variation in modal frequencies was consistent with existing studies reporting their correspondence with phenological phases (Gougherty et al., 2018; Jaeger et al., 2022). For two comparably sized white ash (*Fraxinus americana*), Jaeger et al. (2022) observed a similar seasonal pattern for modal frequencies over one growing season with additional variation in the early spring caused by flowering. Unlike earlier studies, the method used for modal identification in this study also provided information about seasonal variation in other modal properties, including damping ratios, partial mode shapes, and modal force PSDs. The large increase in *ζ*_*i*_ during periods with leaves showed the substantial contribution of aerodynamic drag on leaves towards total damping, and the difference likely explains the larger range for *S*_*ii*_ during period with leaves. Compared to the spring, the smaller change in *f*_*i*_ during the fall could also be explained by phenology. The 0.28 Hz decrease in *f*_*i*_ during the spring was measured alongside an increase in the tree’s mass and height, and the 0.21 Hz increase in *f*_*i*_ during the fall was mostly associated with a decrease in mass. Based on published information (Abrams et al., 1990; Niklas, 1994), the tree’s total dry leaf mass was likely in the low single digits (approximately 2 kg).

Consistent with this study, existing studies also observed a gradual increase in modal frequencies over the summer (Gougherty et al., 2018; Jaeger et al., 2022), and the authors largely attributed the variation to gradual changes in leaf condition caused by aging, stress, or damage. Many have observed a decline in photosynthetic capacity over the summer season (Grassi et al., 2005; Grassi and Magnani, 2005; Wilson et al., 2000), often proportional to drought intensity (Xu and Baldocchi, 2003), and it is reasonable to expect that decreasing total mass from slow drying could explain the gradual increase in modal frequencies. The gradual increase in modal frequencies could also be caused by secondary growth, since the frequency of a cylindrical beam is proportional to its diameter (Gardiner, 1991; Moore and Maguire, 2004). Unfortunately, neither leaf condition nor trunk diameter were monitored over time in this study. Among nine balsam poplars (*Populus balsamifera*) growing on different sites, modal frequencies changed more on some trees than others over the summer (Gougherty et al., 2018), and the differing rates of change suggest site conditions may interact with the underlying process to affect changes in modal frequencies over the summer. The precise cause of the gradual change should be investigated carefully in future studies.

Although some existing studies used 24 h time intervals for estimating modal frequencies (Gougherty et al., 2018; Jaeger et al., 2022), the higher temporal resolution of our estimates obtained from 1 h intervals also revealed other short-term patterns, including local spikes and diurnal trends. Local negative spikes were dispersed throughout the growing season and broadly corresponded with rainfall events, and the mass of intercepted water in the crown likely caused the transient decrease in *f*_*i*_ (Ciruzzi and Loheide, 2021). One clear positive spike in frequency occurred during severe freezing (−10° C) in the early morning on 13 April 2022, and the stiffness of frozen wood likely caused the transient increase in *f*_*i*_ (Granucci et al., 2013).

Diurnally, frequencies varied between a maximum during the night and minimum during the day, possibly caused by a change in the tree’s water content or other exogenous factors. Ciruzzi and Loheide (2019) observed a similar diurnal pattern in the sway period, 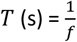, of bigtooth aspen (*Populus grandidentata*) and northern red oak (*Quercus rubra*), and the authors attributed the variation to differences in stored water, inferred from measurements of wet leaf mass per unit area, LMA (g·m^−2^), over 24 hours. They also implausibly suggested the higher moisture content caused an increase in wood stiffness due to turgor pressure, despite the absence of vacuoles from xylem tissue (Evert, 2006) and the higher stiffness of dry wood (Senalik and Farber, 2021). The diurnal variation in modal frequencies could also be explained by changes in dry LMA over daily cycles. Some studies reported LMA increased between 10 and 50% during the day (Poorter et al., 2009), likely due to the accumulation of non-structural carbohydrates or other substances (Bertin et al., 1999; Tardieu et al., 1999). The shrinking and swelling of woody stems over daily intervals from sap movement (Chan et al., 2016; Pastur et al., 2007) could also physically explain the variation. Despite many possible explanations, there is clearly not a convincing account of the reason(s) for daily cycles in modal frequencies, and the many possible explanations should be carefully tested in future studies. Notably, similarly pronounced diurnal patterns in *f*_*i*_ were not observed over a shorter two-week period on a large *Hopea odorata* tree (Burcham and Au, 2022).

The covariation of *f*_*i*_ and *ζ*_*i*_ with *S*_*ii*_ was consistent with earlier findings (Burcham and Au, 2022) and similar reports for built structures (Tamura et al., 1993), and the observations confirmed the broader existence of amplitude dependence in the modal properties of trees. The pronounced influence of leaves on the relationship between modal properties and *S*_*ii*_ was expected, due to the higher drag on leaves, but the observed covariation during seasonal periods without leaves demonstrates the importance of other features and processes in maintaining the relationship. Although much less pronounced on leafless trees, the stratification of modal properties across a range of values at a given amplitude was also consistent with existing observations of built structures (Zhang et al., 2020), and some of the variance was likely caused, at least in part, by changes in environmental conditions (Clinton et al., 2006; Yuen and Kuok, 2010). For example, a small portion of *f*_*i*_ were noticeably lower than others at smaller *S*_*ii*_, and many of such estimates corresponded to wet days with unknown additional mass from intercepted water. Although wet days coincided with many deviations from the general relationship, they did not match all incongruities with the broader trend. When investigating amplitude dependence in the future, it will be important to properly account for changes in modal properties caused by environmental conditions (Zhou and Li, 2021).

The persistence of two close, nearly orthogonal modes was similar to earlier observations of a large tropical tree (Burcham and Au, 2022), and the findings suggest the vibration behavior may be common to more trees across other sites. In built structures, close modes often occur in tall, slender structures with multiple axes of transverse symmetry, and most trees exhibit very similar characteristics. During leafless periods, the orientation of ***φ***_*i*_ may have been explained by the tree’s asymmetrical crown widths with ***φ***_1_ and ***φ***_2_ generally oriented orthogonal to the larger and smaller widths, respectively. The differences in crown size plausibly explain the arrangement of other modal properties observed for the corresponding modes (Kane et al., 2014). While crown architecture may explain the orientation of modes during leafless periods, the orientation of ***φ***_*i*_ appears to be more closely related to the direction of fluid motion during periods with leaves. If the two close modes were caused by nearly axisymmetric conditions, as is the case for built structures, the creation of strong asymmetry from crown pruning or root trenching may remove the conditions leading to the existence of close modes, and the expectation, if confirmed, could provide one basis for tree structural health monitoring.

The uncertainty laws showed that the prevalence of very close modes is a major challenge for modal identification with trees. For built structures, *d*_*i*_ usually exceeds 0.1, but many of the values were much smaller for the green ash. Many of the cases with nearly identical modes corresponded to those with interior angles close to 0° or 180°, and the uncertainty laws do not apply for degenerating cases with two modes roughly aligned in the same direction. In other cases, the algorithm may have converged poorly or suffered from low s/n ratios. For a variety of reasons, nearly identical modes are challenging to address theoretically and computationally, but the use of better sensors with higher sensitivity and lower noise may improve some of the factors contributing to questionable estimates in this study. The sensor (LIS3DSH, STMicroelectronics, Geneva, Switzerland) used in the AL100 has a noise PSD of around 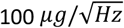, and there are other, similarly affordable MEMS accelerometers available with characteristics better suited to tree vibration monitoring. To improve estimates, the development and testing of alternative sensors with low noise and high sensitivity over the relevant frequency range should be a main priority for future work.

## Conclusions

This study characterized the substantial, often overlapping patterns of variation in modal properties over an entire season for a common deciduous broadleaf tree. Although some changes directly corresponded to other observations in this study (e.g., amplitude, phenology, temperature, rainfall), the causes of other changes were not clear, and it will be important to carefully test some of the possible explanations, such as leaf condition, water movement, and secondary growth, in future work. In agreement with earlier work, the prevalence of close or nearly identical modes appears to be a unique, persistent feature of tree vibration, and the existence of very close modes often limited the number of high-quality estimates, producing a noticeably incomplete description of modal property variation over time during periods with leaves. Given the clear potential for detecting specific environmental changes with tree vibration monitoring, there is a need for more observations of other trees to characterize the prevalence of close, nearly orthogonal modes and determine the causes of natural variability in modal properties over time.

## Data availability statement

The tree movement and wind measurements used in this study were deposited in the Harvard Dataverse at https://doi.org/10.7910/DVN/ZTYPJE.

## Acknowledgments

Colorado State University funded this study. The authors thank Sarah Wilhelm and David Leinbach for their assistance with field instrumentation.

