## Supplementary Materials for "Tracking seasonal variation in modal properties of an open-grown green ash (*Fraxinus pennsylvanica*)"

Table S1 Vegetative phenophase definitions for angiosperm trees

| Phenophase | Definition |
|---|---|
| Breaking leaf buds | One or more breaking leaf buds are visible on the plant. A leaf bud is considered "breaking" once a green leaf tip is visible at the end of the bud, but before the first leaf from the bud has unfolded to expose the leaf stalk (petiole) or leaf base. |
| Increasing leaf size | A majority of leaves on the plant have not yet reached their full size and are still growing larger. Do not include new leaves that continue to emerge at the ends of elongating stems throughout the growing season. |
| Leaves | One or more live, unfolded leaves are visible on the plant. A leaf is considered "unfolded" once its entire length has emerged from a breaking bud so that the leaf stalk (petiole) or leaf base is visible at its point of attachment to the stem. Do not include fully dried or dead leaves. |
| Colored leaves | One or more leaves show some of their typical late-season colors. Do not include small spots of color due to minor leaf damage or dieback on branches. Do not include fully dried or dead leaves that remain on the plant. |
| Falling leaves | One or more leaves are falling or have recently fallen from the plant. |

Denny, E.G., Gerst, K.L., Miller-Rushing, A.J., Tierney, G.L., Crimmins, T.M., Enquist, C.A.F., Guertin, P., Rosemartin, A.H., Schwartz, M.D., Thomas, K.A., Weltzin, J.F., 2014. Standardized phenology monitoring methods to track plant and animal activity for science and resource management applications. Int. J. Biometeorol. 58, 591–601.
